# Learning the architectural features that predict functional similarity of neural networks

**DOI:** 10.1101/2020.04.27.057752

**Authors:** Adam Haber, Elad Schneidman

## Abstract

The mapping of the wiring diagrams of neural circuits promises to allow us to link structure and function of neural networks. Current approaches to analyzing *connectomes* rely mainly on graph-theoretical tools, but these may downplay the complex nonlinear dynamics of single neurons and networks, and the way networks respond to their inputs. Here, we measure the functional similarity of simulated networks of neurons, by quantifying the similitude of their spiking patterns in response to the same stimuli. We find that common graph theory metrics convey little information about the similarity of networks’ responses. Instead, we learn a functional metric between networks based on their synaptic differences, and show that it accurately predicts the similarity of novel networks, for a wide range of stimuli. We then show that a sparse set of architectural features - the sum of synaptic inputs that each neuron receives and the sum of each neuron’s synaptic outputs - predicts the functional similarity of networks of up to 100 cells, with high accuracy. We thus suggest new architectural design principles that shape the function of neural networks, which conform with experimental evidence of homeostatic mechanisms.

## INTRODUCTION

Many biological systems can be described as networks of interacting elements where the function of the system is determined by the nature of individual elements, the type of interactions between them, and the emerging individual and collective behavior or phenotype. Mapping the relation between the structure and function of such networks is a key goal in many areas of biology, and beyond. Because the number of possible architectures is combinatorial in the size of the network, analyzing and understanding the design and function of biological networks hinges on finding simplifying principles. Functional ‘design principles’ have been suggested to include robustness to noise [1–3], resilience to attack [4], controllability [5], efficiency [6, 7], criticality [8, 9], and learnability [10, 11]. Structural design principles have implied the nature of network growth [12], use of small sub-network motifs [13], modular organization and power-law scaling [14, 15], sparseness of activity [16], random connectivity [17, 18], and centrality or percolation properties of networks [19, 20]. However, how these functional and structural design principles relate to one another is not immediately clear [21].

The reconstruction of the detailed wiring diagrams of full neural circuits at single cell resolution [22–26], would enable direct exploration and characterization of the architectural design of neural modules and even whole brains [27]. Importantly, very different connectivity structures may give rise to very similar function [28]. Thus, the ability to record the joint activity patterns of large populations of neurons [29, 30] whose *connectome* has been reconstructed, is crucial for linking of neural network structure and function [31–34] - which would be central to our understanding of development, coding, plasticity, and learning in biological neural networks.

Understanding the relations between network topology and the activity of networks of neurons requires ways to measure both the functional similarity of networks and their architectural similarity, and to map the relations between these two (potentially very different) metrics. It is not obvious what is the correct measure for either or how we may extend tools from graph and network theories [12, 15, 35]. For limited classes of networks and of interacting elements, links between the topology of a network and the nature of its dynamics have been elucidated, such as the number of fixed points or classes of attractor dynamics and their stability [36–40]. Towards the study of real neural networks, we present here a general framework for linking the topology of networks and their population spiking patterns for arbitrary classes of network architectures, in response to a wide range of stimuli.

Rather than assuming or guessing which structural metric should be used to compare networks - we learn a functional similarity metric directly from the networks. We thus characterize the space of neural networks in terms of their function and then seek the architectural principles that govern the organization of that space. We develop this framework and validate it by studying simulated networks of spiking neurons, where we have complete control over all parameters, no limits on experimental design and length, and the ‘ground truth’ is known. We show that we can *learn* the informative structural features that shape a network’s function, and that a structural metric, based on these features, significantly outperforms a wide range of graph-theoretical measures in predicting the functional similarity of neural circuits. We then show that the informative structural features that we identified for small networks are highly informative also for networks of 100 neurons - suggesting them as a general principle for the comparison of networks of neurons.

## RESULTS

To study the relation between structure and function in networks of neurons, we simulated the responses of thousands of small networks of spiking neurons to a wide range of stimuli (Fig. 1a). Direct characterization of the space of all network architectures is impossible for large networks, since for a group of *N* neurons there are 2^*N*(*N*−1)^ different directed graphs of interactions (topologies); considering neurons of different types or diverse strengths of synaptic connections, would make this number considerably larger. We therefore start by seeking the principles that govern ensembles of small networks and later increase their size (Fig. 1b). The first two ensembles are the exhaustive sets of all 4,096 topologies of networks of 4 all excitatory or all inhibitory leaky integrate and fire neurons, with all synapses having the same strength (we simulated both current and conductance based models for the neurons and both alpha and delta activation functions for the synapses - see SI). The next ensemble was comprised of networks of 15 excitatory and inhibitory neurons, with ~ 20% inhibitory cells [41], where synaptic strengths were drawn from a log-normal distribution [42]. Even for 15 neurons there are ~ 10^63^ directed topologies, and so we used a random sample of 10,000 of networks of each size, and rely on cross-validation on new networks to verify our results and models (see Methods).

**Figure 1.**
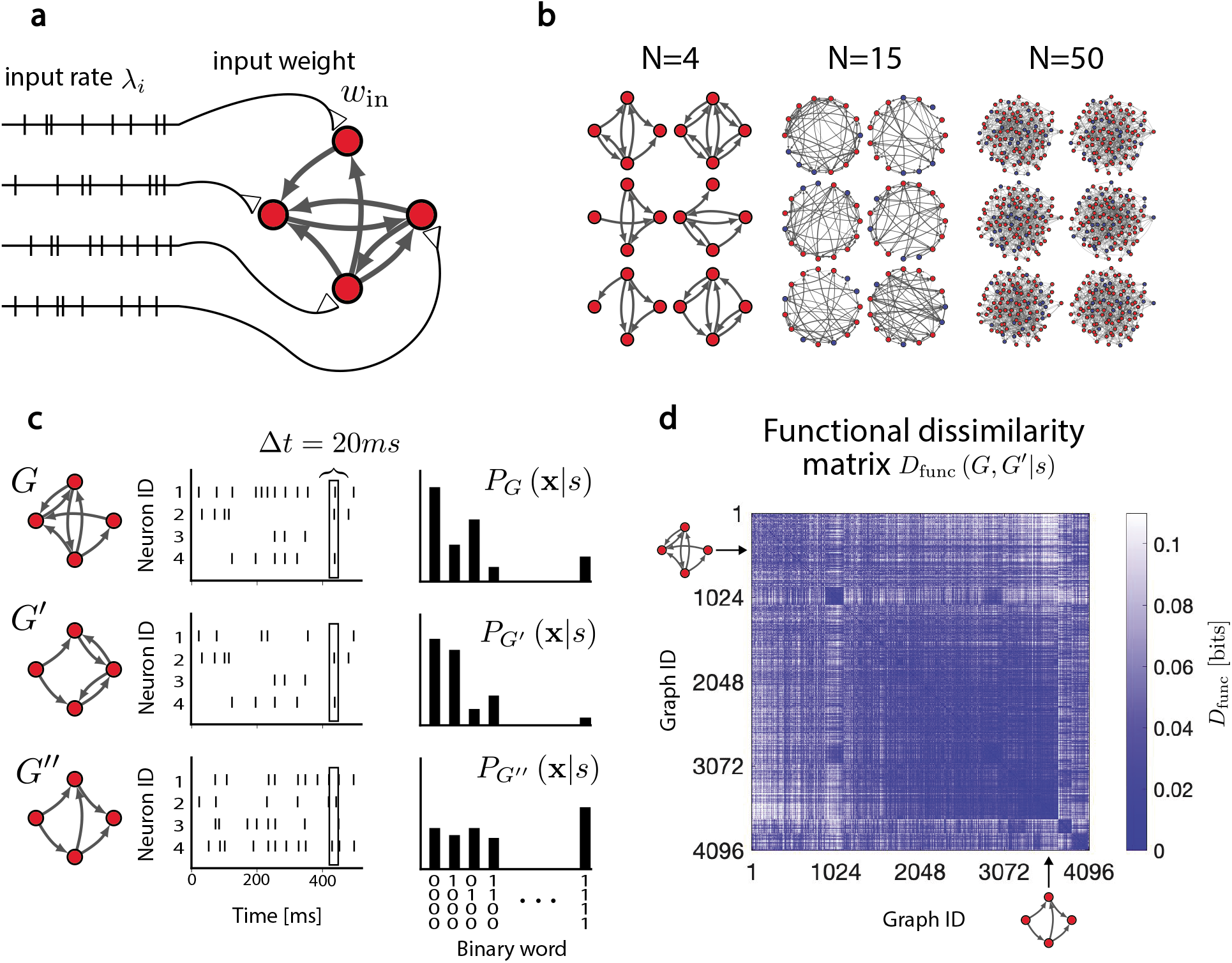
Simulating the responses of ensembles of networks to the same stimuli and computing functional similarity between networks. **(a)** Each of the *N* neurons in the simulated networks received as an input a Poisson spike train with rate *λ_i_*; corresponding neurons in all networks in the ensemble received the same exact i-th spike train as input. **(b)** Examples of the simulated networks: Networks of 4 neurons with identical excitatory synapses, networks of 15 excitatory and inhibitory neurons where the synaptic weights were drawn from a log-normal distribution, and networks of 50 excitatory and inhibitory neurons with log-normal distributed synapses. **(c)** Examples of segments of the spike train responses of different networks (left) that were presented with the same stimulus. The activity of the neurons was discretized into bins of length Δ*t* = 20*ms*, and binarized (middle; see text). The different responses of the networks result in different distributions of population activity patterns (right). **(d)** The functional dissimilarity matrix *D_func_* (*G*, *G′|s*) for networks of size *N* = 4. The normalized stimulus strength was *η =* 1.5 (see Methods), which resulted in firing rates ranging between 10Hz-20Hz over all networks in the ensemble. The structure of this matrix implies a low dimension organization of the functional space of networks (see text).

To map the functional similarity between networks, we simulated their responses to the same stimulus *s* and compared their respective population activity patterns (Fig. 1c). We denote each network by a weighted connectivity graph or the matrix of synaptic connection *G*, where *G_kl_* is the strength of the synapse from neuron *k* to l. The external stimulus s to the networks was defined individually for each neuron, such that the i-th neuron in each network received as an input a 30 seconds Poisson-distributed spike train with a rate *l_i_*, weighted by an input synaptic weight, *w_input_* (Fig. 1a). While the rate *l_i_* of inputs to all neurons was identical, each neuron in the network received its own realization of incoming spikes with that rate (where corresponding neurons in different networks received the *exact* same input patterns, i.e. same realization). We explored the responses of the networks to a wide range of stimuli (SI and Fig. S1) and focus henceforth on stimulus parameters and synaptic strength values for which the networks’ behavior was not pathological, i.e. epileptic or completely silent. The length of the stimulus was chosen to give an adequate sample of the responses that this class of stimuli would elicit. To disentangle the effects of architectural differences between networks from the effects of the initial conditions of networks on their responses - all networks were started with the same initial conditions, and all network measures and similarity measures between networks are the average over many different random initializations. We discretized the spiking patterns of the neurons in the network in response to the stimulus, into small temporal bins of size Δ*t* = 20 ms, such that the activity of the network in each time bin is given by a binary vector **x** = (*x*_1_,…,*x*_*n*_), where *x_k_ =* 1 if neuron *k* spiked in that bin and 0 otherwise (Fig. 1c, middle). We then summarized the response of the network whose synaptic connections are given by *G*, to stimulus *s* by the distribution of population activity patterns over the whole length of the stimulus (Fig. 1c, right), which we denote by *P_G_*(**x**|*s*). For small networks we estimated these distributions by direct sampling, whereas for large ones we fitted a pairwise maximum entropy model for the population activity (see Methods). Using different temporal bin sizes gave similar results for the analyses that follow (SI and Fig. S2).

We quantified the functional similarity of pairs of networks whose synaptic weights are given by *G* and *G*′, by the overlap of their population responses, conditioned on the stimulus *s*:

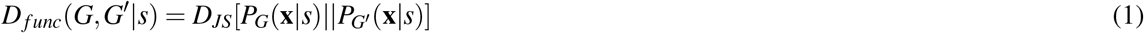

where *D_JS_* is the Jensen-Shannon divergence, which measures the differences between distributions in bits (see Methods). Notably, this comparison of the networks’ population vocabulary to a particular stimulus or class of stimuli is more general than overlap measures based only on individual firing rates, as it also considers differences in correlation structure between neurons. As this does not take into account the temporal structure of the responses, we also compared networks based on the similarity of the post-stimulus-time histogram (PSTH) of the corresponding cells in the two networks to the same stimulus, *D_PSTH_* (see SI). We found that *D_PSTH_* was strongly correlated with *D_func_* (see SI and Fig. S3), and we therefore focused on the latter for the rest of the analyses.

We computed the functional dissimilarity between all networks in our ensembles. Figure 1d shows a typical example of the resulting matrix of dissimilarity between pairs of networks, *D_func_*(*G. G′ |s*), for s with a normalized input rate of *η =* 1.5 and *w_input_ =* 20*pA* (which is equivalent to a total rate of incoming spikes to each neuron of 8Hz, see Methods). The structure of the matrix reflects a low dimensional organization of the space of networks based on their response properties - evident by the spectrum of the eigenvalues of the matrix, which decayed significantly faster than shuffled controls (SI and Fig. S4). We then asked what are the structural properties of the networks that underlie this functional organization.

### Common structural metrics fail to capture the functional similarity of networks of neurons

Given the plethora of graph theory measures of similarity, it might seem that a smart choice of one such measure might be sufficient to predict the functional similarity of neural networks. The simplest intuitive way to compare the topology of networks of the same size is by counting the fraction of links or synapses they share. For weighted directed graphs, this can be interpreted in different ways, and we consider here two options: First, the Graph Edit or Hamming distance between *G* and *G*′, 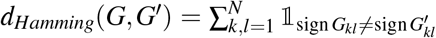, (where 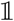 is an indicator function), which compares the *type* of connections between neurons (inhibitory, excitatory, or absent). Second, the *L*_2_ norm or Euclidean distance between the corresponding synapses in the networks, 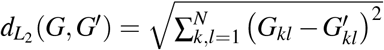, where *G_kl_* is the synapse from neuron *k* to *l* in network G. Both of these metrics were poor predictors of the functional similarity of networks (Fig. 2b,f). In particular, while low Hamming or Euclidean distance did imply low *D_func_*, medium and high values of Hamming or Euclidean distance were practically non-informative of functional similarity.

**Figure 2.**
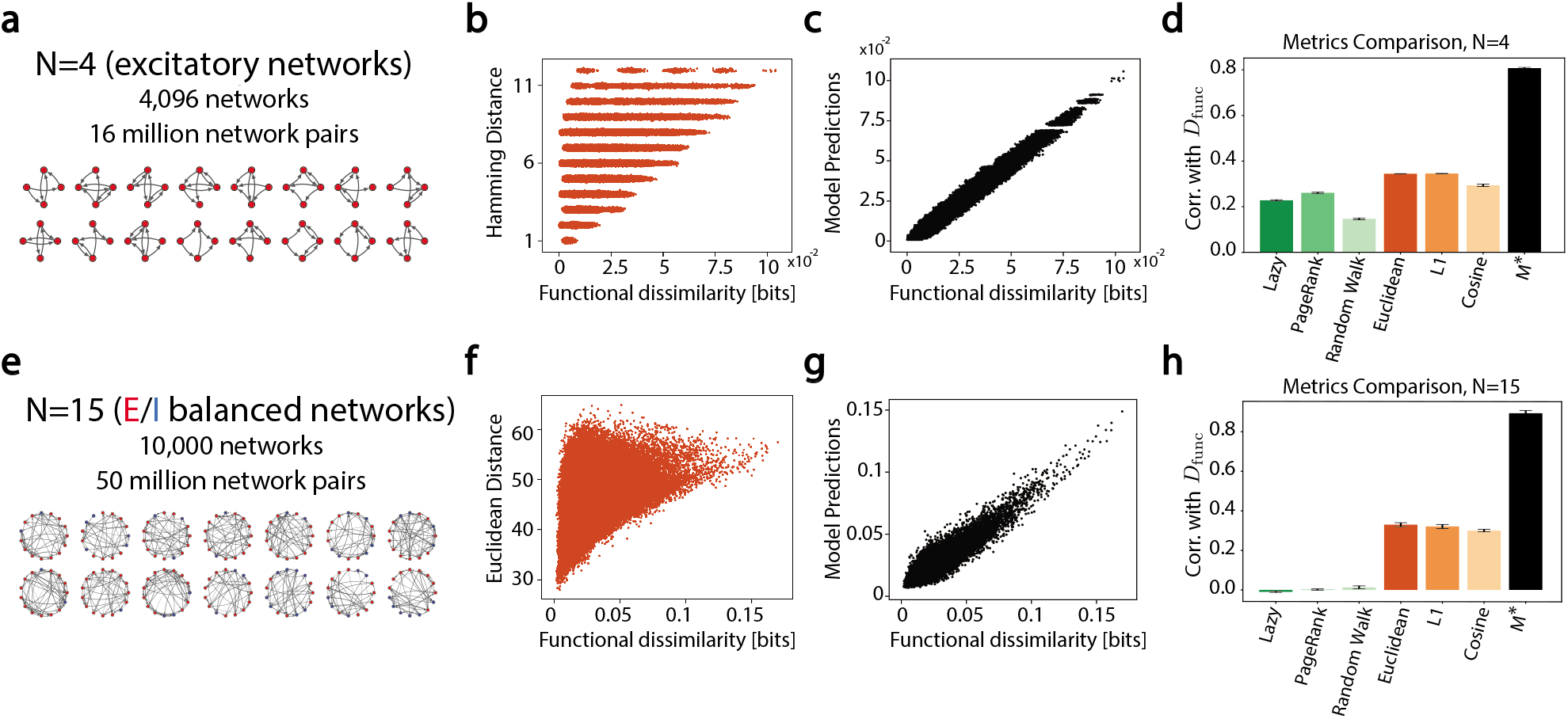
Common structural metrics fail to predict the functional similarity between networks of neurons, whereas a learned bi-linear metric succeeds. **(a)** The first simulated ensemble, consisting of 4096 networks of 4 neurons with constant weights. **(b)** Hamming distances between all pairs of 4096 networks of size *N =* 4 are plotted against the computed *D_func_* between networks (y-values are slightly jittered to show density of points). We note the points on the upper-left part correspond to pairs of networks that have almost no overlap in terms of synaptic connections, yet are functionally very similar. **(c)** Predictions of the bi-linear model on held out test data, i.e. networks that were not used in finding *M*\*. **(d)** We compare the prediction accuracy of different structural metrics for networks of size *N =* 4, by computing their mean correlation with the functional similarity of the networks (averaged over 30 different initial conditions). Our learned model (black) outperform multiple graph-based models (green) and vector-based ones (orange). Error bars represent 1 standard deviation. **(e)** The second simulated ensemble, consisting of 10000 networks of 15 neurons with both excitatory and inhibitory weights. **(f-h)** Same as (b-d), but for the *N* = 15 ensemble. Since synaptic weights in this ensemble were continuous, the Euclidean distance between synaptic weights was used instead of Hamming (see main text).

We examined a wide variety of other structural metrics, including ones that treat networks as vectors of synaptic weights and compares them as vectors in *R*^*N*(*N*−1)^, as well as explicit metrics of graphs, such as the spectral distances between directed graph Laplacians. All structural metrics in our comparisons gave poor results, for both *N* = 4 and *N* = 15 - as reflected by the green and orange bars in Fig. 2d,h (a larger set of failed metrics can be found in SI and Fig. S5). We therefore asked whether instead of trying common similarity measures, we could learn a metric directly from the data.

### Learning to predict the functional similarity of networks from their synaptic connections

Metric learning approaches vary significantly in their assumptions and applications [43], and have been used to model perceptual distances [44, 45], and to study the structure of neural codes [46–48]. Here, we asked how well *D_func_* between networks’ responses to the same stimulus s can be approximated by a bi-linear function of the synaptic differences between the networks, which we quantify as the difference of their connectivity matrices Δ*G_kl_ = G_kl_ − G′_kl_*. To simplify mathematical notation, we denote by 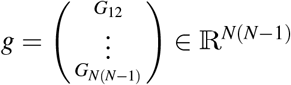 the vector corresponding to the “flattened” representation of the matrix G, without its diagonal elements (which are all zero), and so Δ*g* = *g* − *g*′ is the vector of synaptic differences between two networks. The problem of finding the optimal bi-linear function is then translated into seeking the optimal matrix *M*\*(*s*), which obeys

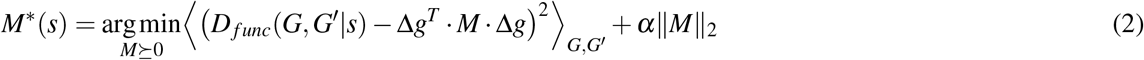

where *M* is a *N*(*N* − 1) × *N*(*N* − 1) positive semi-definite matrix (so defined in order for the distances between networks to be non-negative), known as the Mahalanobis matrix [43]. The regularization term and its control parameter *α* were chosen by cross-validation (SI and Fig. S6). Fortunately, *M*\*(*s*) we seek is the solution to a convex constrained optimization problem, which is therefore guaranteed to be the global optimum. We explicitly note the dependence of *M*\* on *s*, and stress that different stimuli might require different metrics, which we investigate below.

To find *M*\*(*s*), we randomly split the networks in the ensemble into a training set and a test set (75% and 25% of the networks, respectively), and use the train set to find *M*\*(*s*) using conjugate gradient descent on the manifold of positive semi-definite matrices (see Methods). To assess how well this learned measure captures the functional dissimilarity between networks, we use it to predict the pairwise distances between all pairs of networks in our held-out test set and compare these predictions to the empirical *D_func_* values (Fig. 2c,g). We find that our model captures functional dissimilarity significantly better than all other metrics (Fig. 2d,h), for both networks of size *N* = 4 and *N* = 15. We emphasize that the predictive accuracy of *M*\* stems from its ability to capture the geometry of the functional space of networks, and not from its computational expressive power (see SI and Fig. S7).

The sparse structure of the Mahalanobis matrix *M*\* for networks of 4 excitatory neurons (Fig. 3a), reflects the architectural features that the model relies on. To uncover what these features are, we used the fact that *M*\* is a positive semi-definite matrix that can be decomposed into its *Cholesky factor, M*\* = *RR^T^*, such that *R* is a lower-triangular matrix. Importantly, we can use *R* to find a decomposition of *M*\* that is easier to interpret: *M*\* = *LL^T^*, with *L = RU* and *U* is unitary matrix that would make *L* as sparse as possible (see Methods). Using this sparse decomposition, the distance between networks can be rewritten as 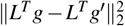, which means that *L^T^* implements a linear transformation that represents each network using a set of structural features. In this view, our model measures the squared Euclidean distance between networks in the feature space induced by *L^T^* (which interestingly is consistent with *DJS* being a squared metric). The interpretation of the structural features extracted by *L* (Fig. 3b), is facilitated by the fact that each column of *L* defines a linear combination of synaptic weights. For the ensemble of excitatory networks of size *N =* 4, a dominant type of structural features stands out - one of the form ∑*_l_ G_lk_*, or the *sum of synaptic inputs* to the k-th neuron (this corresponds to the *indegree* in unweighted networks). Thus, for example, the left column of *L* in Figure 3b is akin to the sum of synapses going into the third neuron. We identified two other types of features when using other stimuli and for networks with inhibitory neurons: the sum of outgoing synapses from a neuron, ∑*_k_G_lk_*, and the sum of synaptic weights along loops of size 2, *G_kl_* + *G_lk_;* the last type was dominant for example for the ensemble of all inhibitory networks of size *N* = 4, as shown in Fig. 3d-e. Since each column of *L* corresponds to a single structural feature, we further asked how many features are needed to accurately approximate the functional dissimilarity. We sorted the columns of *L* by their norms (as vectors), and approximated *M*\* using the k-th highest norms columns, for different values of k. We found that for both networks of size *N* = 4 and *N* = 15, the number of the required structural features was small, and close to the number of *neurons* rather than the number of *synapses* (Fig. 3c).

**Figure 3.**
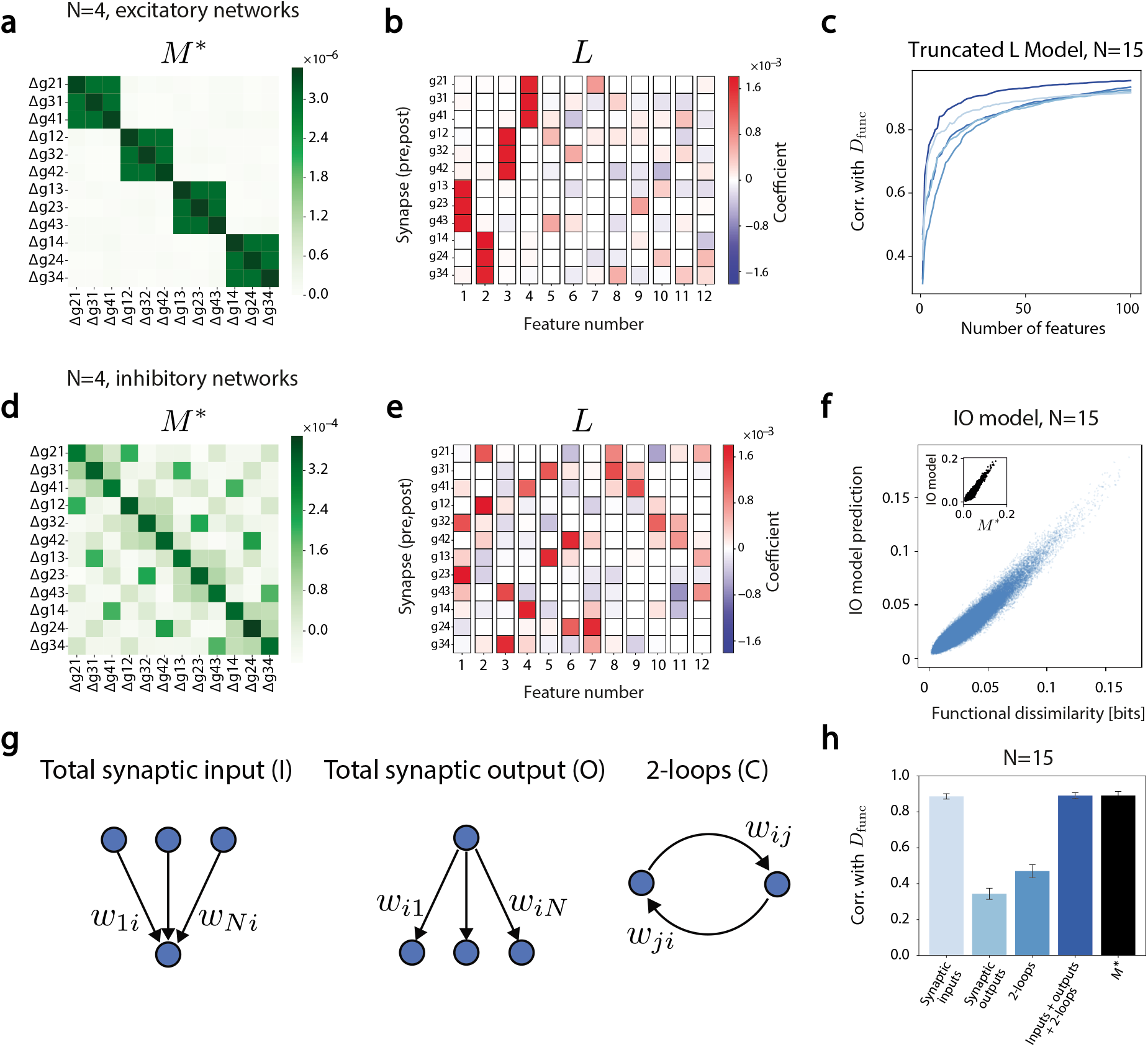
Identifying the informative structural features that govern our learned model of network similarity. **(a)** The optimal Mahalanobis matrix *M*\* for *N* = 4 networks of excitatory neurons. Each entry in the matrix is the weight assigned to a pairwise term in the bi-linear function. **(b)** The sparsest *L* matrix from Cholesky factor-based decomposition of *M*\* from (a). Each column represent a single structural feature. Matrix entries correspond to the weights of the different synapses in each feature. First 4 features correspond to the total synaptic inputs to each of the neurons; Columns are sorted by their Euclidean norm. **(c)** Accuracy of the model when using only the k most important features (largest Euclidean norm), for networks of *N*=15 and different stimuli. **(d,e)** Same as (a,b) but for networks of 4 inhibitory neurons. Here loops of length 2 emerge as the dominant structural features. **(f)** A model based only on the sum of synaptic inputs to each neurons and the sum of synaptic outputs of each neuron, predicts the functional similarity of networks of *N* = 15 neurons, as well as the full model using *M*\*; inset shows the high correlation between the bi-linear model and the feature-based model. **(g)** An illustration of the informative structural features. **(h)** Prediction accuracy (mean pearson correlation coefficient) using different subsets of structural features.

Based on these results, we fitted anew model that uses just the dominant structural features of *L* to approximate *D_func_*(*G*, *G*′|*s*):

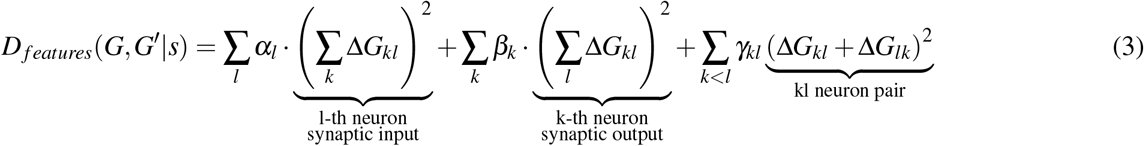

where *α_l_. β_k_. γ_kl_* were learned from the data to minimize the mean squared error between the two measures. For many stimuli, the features-based approximation was close to the full learned matrix *M*\*. Furthermore, an even simpler class of models that relied just on the first two terms of eq. (3), namely the set of sums of synaptic inputs and sums synaptic outputs of each of the neurons - was nearly as good, as shown in Fig. 3f.

### Scaling and universality of the functional space of networks for different classes of stimuli

The results above showed the success of learning a feature based metric for networks for a specific set of stimuli, where all neurons received inputs from different realizations of Poisson distributed spike trains with the same rate. Next, we asked how the functional similarity between networks depends on the nature of the stimulus used, and in particular, whether the functional metric we learned for one class of stimuli would generalize to other stimuli. We therefore repeated the mapping of functional distances between networks, *D_func_*, for a wide range of Poisson-inputs with different mean values - from weak stimuli that elicited almost no responses, to very strong stimuli that elicited very high firing rates (SI and Fig. S1), and from uncorrelated inputs to the different neurons to highly correlated ones (Figure 4a). We found that the distances between networks were highly correlated for different stimuli: Figure 4b shows an example of the tight correlation of *D_func_*(*G*, *G′|s*) for two different stimuli, over all pairs of networks in the ensemble. Figure 4c shows the strong correlation between the distances of networks for many different pairs of stimuli, which were high and significant for all stimulus classes we tested (see SI and Fig. S8 for the correlation values of the dissimilarity of all the stimuli we used throughout the main text). This implies that on average, changing the stimulus changes the dissimilarity between networks in a way that preserves the relations between their relative distances: networks that are relatively close (far) under one stimulus *tend* to be relatively close (far) under a different stimulus. Figure 4d shows this explicitly by using 2D embedding of networks based on their functional distances, using an example of the same randomly selected 4 networks under 5 different stimuli - reflecting that the structure of relations between networks is preserved, while the overall map of distances may stretch or shrink.

**Figure 4.**
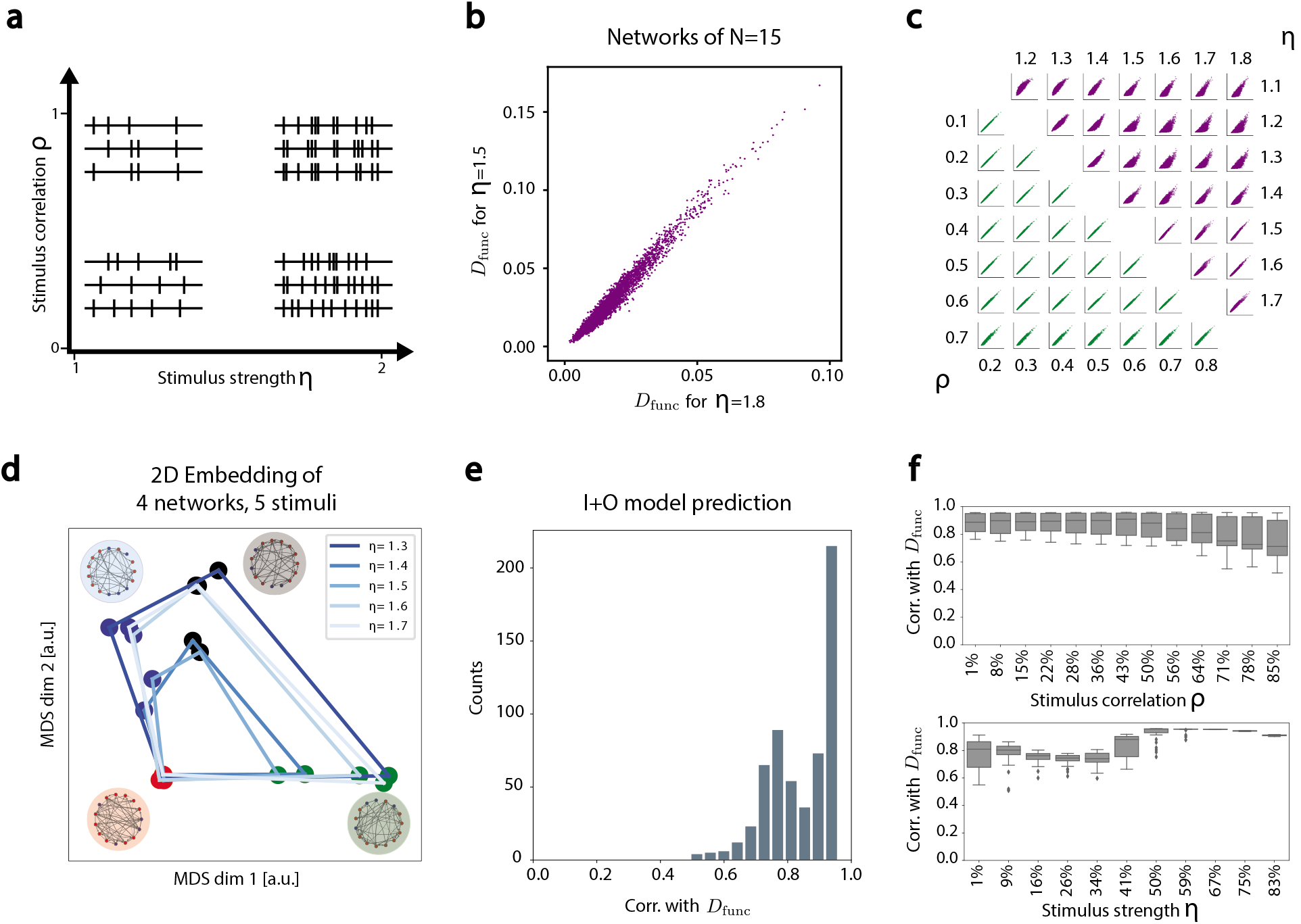
The structure of the map of functional similarities between networks is conserved across
stimuli. **(a)** Illustration of the space of different stimuli for which we computed the pairwise similarity between networks. The set of stimuli explored was parametrically characterized by the average rate of the external Poisson input to the neurons, *η*, and the correlation between different inputs to different neurons, *ρ*. **(b)** An example of the correspondence of *D_func_* for 1 million pairs of networks in the ensemble of networks of size *N* = 15, is shown for two different stimuli; every dot represents one pair of networks. **(c)** Same as (b), but for many pairs of stimuli, from the set space described in (a). The correlations between the distances between all pairs of networks are shown for the whole range of r and h values. **(d)** 5 overlaid 2D MDS embeddings of 4 example networks of size *N* = 15, based on the functional dissimilarity between them. The networks in each case are the same, and are marked by a dot of the same color. The colors of the lines between nodes denotes which of the 5 stimuli this embedding relates to. Overlaying was done by anchoring the network marked by the red dot. Overlaid maps show the geometric organization is preserved across stimuli space. **(e)** Pearson correlation between the computed functional similarity *D_func_* between networks and predictions of the models based on the sum of synaptic inputs and sum of synaptic outputs of each neuron, for 900 different combinations of (*η,ρ*) for networks of size *N* = 15. **(f)** The interquartile ranges of the data shown in (e), is shown by aggregating the values shown in (e) over *η* (up) and *ρ* (bottom). Percentage values of each bar denote the parameter value between lowest strength/correlation (0%) and highest ones (100%).

The approximate scaling of the map of functional distances with stimulus strength suggests that the structure of the Mahalanobnis matrix *M*\* should also remain stable, up to a stimulus-dependent multiplicative factor. Figure 4e-f show the accuracy of the models based on the sum of synaptic inputs and sum of synaptic outputs, across the space of stimuli we explored. Indeed, for the vast majority of stimulus parameters, the prediction of our model and the empirical *D_func_* values are highly correlated. Interestingly, the relative importance of the total synaptic inputs and total synaptic outputs changes across stimulus space, and a transition from the domination of the total synaptic output values to the total synaptic inputs occurs as the stimuli become stronger (SI and Fig. S9).

### Predicting the similarity of large networks from their structural features

Finally, we asked whether the structural features we identified for small networks generalize to large ones. Notably, for networks of 100 neurons there are 2^9900^ possible unsigned topologies and 2^100^ possible activity patterns of the network. Thus, whereas mapping all networks was intractable already for 15 neurons, here even measuring the similarity of networks using the overlap of their sampled response distributions becomes impractical. We therefore fit second order maximum entropy models for each network’s activity (see Methods), and use them to measure the functional dissimilarity between networks G and *G*′ by the average of the loglikelihood ratio of the spiking response of G under a model that was learned from the spiking responses of *G*′, and vice versa: 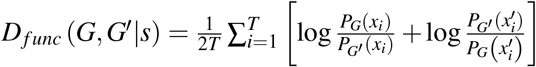, where 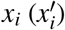 denotes the i-th activity pattern of network *G* (*G*′); This measure converges to the symmetric KL divergence between conditional response distributions, as the number of samples increases.

We used randomly sampled networks of 50 and of 100 neurons, and created 300 variants for each one of these origin networks, by manipulating their connectivity graphs (Fig. 5a): (1) 100 variants in which the sum of synaptic inputs to each neuron was preserved, but the incoming synapses to each neuron were assigned to a random neuron in the network (shuffle inputs). (2) 100 variants that preserved the sum of synaptic outputs of each neuron, but the outgoing synapses of a each neuron were randomly connected to the other neurons (shuffling the outputs). (3) 100 variants in which all synapses were randomly shuffled, regardless of their presynaptic or postsynaptic neuron. As before, all networks were presented with the same stimulus, and had identical initial conditions. Figure 5b-c show the 2D MDS embeddings of all the variant networks of one original network of 50 and of 100 neurons, based on their functional dissimilarity matrix - where closer points represent networks that have a similar conditional response distributions. Figure 5d shows for 50 randomly chosen original networks (with N=100 neurons) that variant networks with the same total synaptic inputs to each neuron (input shuffle) were functionally more similar (smaller *D_func_*) to their original network compared to networks with the same total synaptic outputs for each neurons, which in turn were more similar than networks where all synapses of the original network were shuffled. Figure 5e shows this is the case for all networks we tested, and that this result is consistent under different stimuli. Learning a functional metric as in eq. 2 and the decomposition of such matrices for large networks is impractical, since *M*\* scales as *O*(*N*^4^). Instead, we asked whether the architectural features we identified for small networks would generalize to large ones. We found that using the features based model (eq. 3) we can accurately predict functional dissimilarity between networks of 50 (Figure 5f).

**Figure 5.**
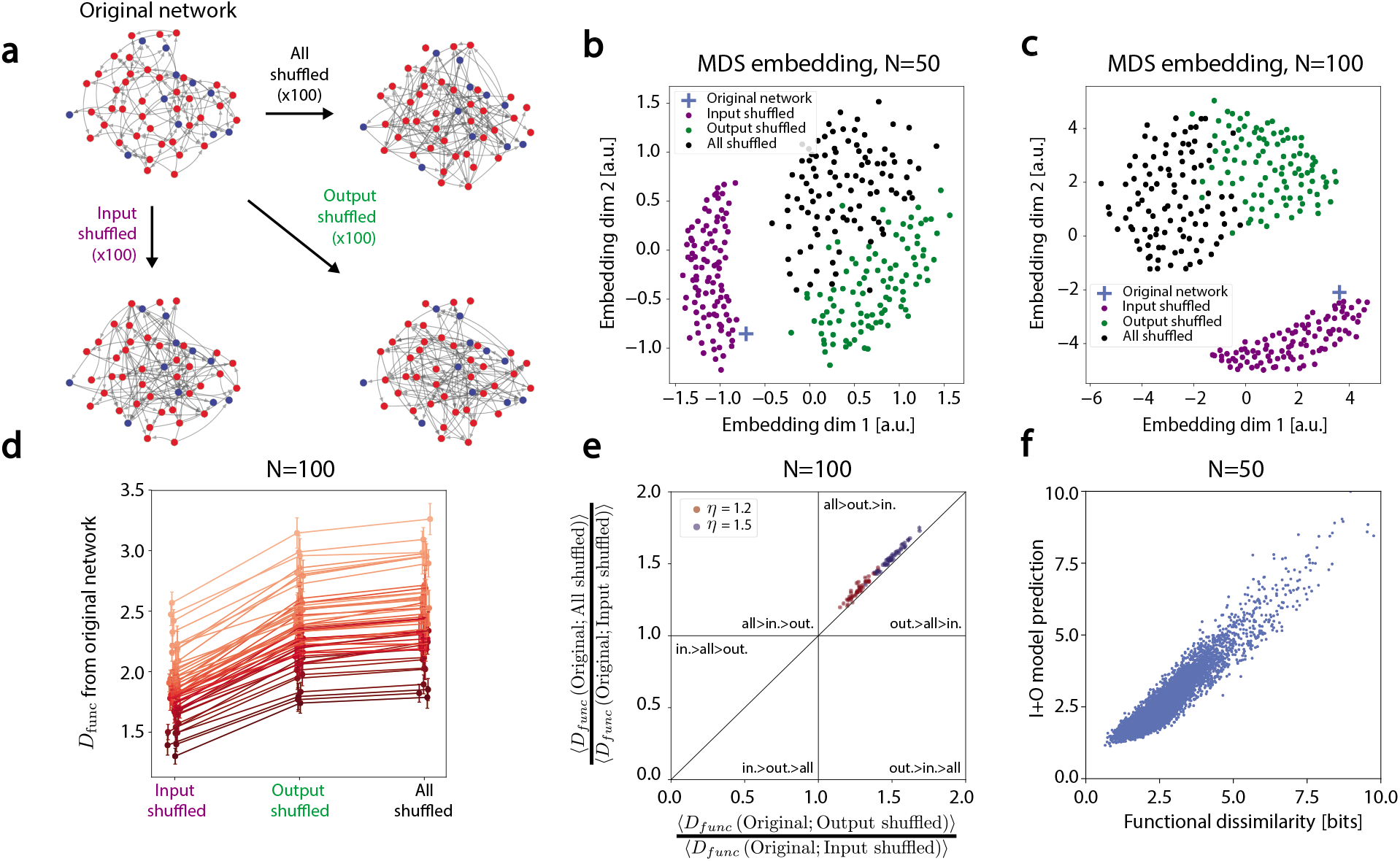
The architectural features identified in small networks predict functional similarity of networks of 50 and of 100 neurons. **(a)** We pick 100 randomly connected networks of size *N* = 50 and *N* = 100, and use each of these networks as a ‘template’. From each such original network, we created 100 variants preserving the sum of synaptic inputs to each neuron, by shuffling the source of the synapses into each neuron (red), 100 variants preserving the sum of synaptic outputs of each neuron, by shuffling outgoing synapses (green), and 100 variants where all synapses shuffled (black). **(b)** A 2D MDS embedding of one original network of 50 neurons and its 300 variants, based on the their log-likelihood based functional similarity values. **(c)** Same as (b) but for a template network of 100 neurons. **(d)** *D_func_* between each template network of 100 neurons and its 3 types of shuffled variants. Each of the 50 lines using different shades of red connects the values of the mean *D_func_* between one original template network and its 100 shuffled variants of each type. Errorbars represent 1 standard deviation over the set of variants. (Small random horizontal jitter was added to the points for clarity). **(e)** For each of the original network, we show the ratio between its average *D_func_* to all the output shuffle variants against the average of its *D_func_* to the all shuffle variants, normalizing both values for each network by its average *D_func_* to input shuffled variants. The colored dots mark the values for the networks under two different stimuli, showing that all 100 original networks reside in the part of the “phase space”, where their input shuffled networks are closer to the original network than the output shuffled ones, which are closer than the all shuffled ones. (The relations between variants in other parts of the phase space are labeled in each of the corresponding parts of the figure). **(f)** Predicting functional dissimilarity, *D_func_* between randomly sampled pairs of networks of 50 neurons using the feature-based model (eq. 3) - showing high correlation with R=0.94.

## DISCUSSION

Measuring the functional similarity of small neural networks in terms of their population spiking patterns, allowed us to learn a structural metric on the space of networks that predicted functional similarity with high accuracy. Our learned metric outperformed a wide range of commonly used graph-theory metrics, by very large margins. We then used the learned metric to identify features of the topology of networks that govern their function. The total synaptic sum of inputs and total synaptic sum of outputs emerged as the key architectural features in small networks, and generalized to predict the functional similarity of networks of up to 100 neurons. We further found that the ‘geometrical organization’ of distances between networks is highly correlated over a wide range of stimulus strengths and correlations.

Our results imply that within the space of networks there are subspaces or manifolds that retain similar functional properties, and that these manifolds are the ones which contain networks that have the same sum of synaptic inputs and sum of synaptic outputs. A network that would change its synaptic connections along a trajectory contained within such a manifold, could therefore be very different structurally, without changing its function. This implies a ‘neutral evolution’ path for neural networks’ organization and learning dynamics [49]. Interestingly, our results conform with the synaptic homeostatic mechanisms that have been extensively studied both experimentally and theoretically [50, 51]. In particular, our model predicts that homeostatic mechanisms that redistribute synaptic weights but preserve the total synaptic inputs to a neuron or its outputs - may not only shape the computational properties of single cells, but play a crucial role in learning, plasticity, and development at the level of the network.

While there are many potential extensions of our work in terms of more biologically-realistic neurons and neuronal classes, we especially note that our analysis has focused on the cases where all neurons in a network receive external stimuli that have the same statistical properties. Extending our work to the case of arbitrary stimuli, such as different input rates to each of the neurons, or cases with higher order correlations among inputs, would require learning a joint metric for stimulus space and the space of network architectures. These are likely to reveal new, more intricate design features of neural circuits. In addition, while our analysis has focused on stimulus values that did not result in pathological behavior of the network (silence or epilepsy), the framework we introduced may be applied to identify which network topologies would be susceptible to such events.

The metric we have learned gave accurate results by relying only on the differences between corresponding synapses of two networks. To explain the small residual part of the functional similarity between networks that our metric did not account for would require to go beyond pairwise relations between synapses. Such extensions could elucidate the functional importance of longer loops and global structural properties, which our current model cannot account for. Moreover, our metric also implies the functional difference between networks would be ‘translation invariant’ in *δG*, i.e. *D_func_*(*G*, *G′|s*) = *D_func_*(*G* + *δG. G′* + *δG|s*) for any *δG*. This is unlikely to be true in general, especially for large dG. We thus expect that refinements of the approach we presented here, such as learning different local metrics, would be important for analyzing larger networks and real networks, and are likely reveal additional design principles.

The growing abundance of connectomics data, would make it imperative to combine theoretically grounded computational frameworks and experimental measurements in understanding networks’ structure and function. The approach we presented here is a step towards building such a framework. While we relied on simulated networks, our results suggest testable predictions about fundamental design principles of real neural networks. Ultimately, this kind of approach would be extendable to asking how to design neural circuits that would perform a desired function, or how to interface and control neural circuits in the brain. Finally, we note that the framework we presented here would be immediately extendable to studying other biological and non-biological networks.

## MATERIALS AND METHODS

### Simulating neural networks

All networks were simulated using the NEST simulator for spiking neural network models [52]. Networks of 4 neurons-either all excitatory or all inhibitory ones - were simulated using the following parameters for an integrate and fire neuron model (which are the default Integrate and Fire model parameters in NEST; “iaf psc alpha”): resting membrane potential *E_L_* = −70*mV*; membrane capacity *C_m_* = 250*pF*; membrane time constant *τ_m_* = 10*ms*; refractory period *τ_ref_* = 2*ms*; spike threshold *V_th_* = −55*mV*; reset potential *V_reset_* = −70*mV*; rise time of the synaptic alpha function *τ_syn_* = 2*ms*. Networks of 15 excitatory and inhibitory neurons were simulated using the biophysical parameters taken from [41]: *E_L_* = 0*mV*; *C_m_* = 250*pF*; *τ_m_* = 20*ms*; *τ_ref_* = 2*ms*; *V_th_* = 20*mV*; *V_reset_* = 0*mV*; *τ_syn_* = 0.5*ms*. Networks of 50 and of 100 neurons were simulated using the same biophysical parameters as for networks of 15 neurons.

Mean synaptic weight were chosen such that an input spike to a neuron results in a 0.1mV increase in its membrane potential. Other mean synaptic weights (ranging from 0.05mV to 0.5mV increase per spike) were also considered and gave similar results.

The external stimulus to each neuron was a sequence of spikes drawn from a Poisson distribution with rate *λ_i_*, with a fixed synaptic strength (*w_inpu_ =* 20pA). The rate was chosen such that *λ_i_ = η · λ_th_*, where *λ_th_* is the external rate that would fix the membrane potential of the receiving neuron around its threshold and *η* is the the ratio between the external rate and the threshold rate, as in [41]. Stimulus values ranged from 5000Hz to 15000Hz (thus simulating the inputs from hundreds to thousands of external neurons), yielding average firing rates in the simulated neurons that range from 1Hz to 50Hz (see SI and Fig. S1). Correlated stimuli were generated using a multiple interaction process [53]; we considered stimulus correlations ranging from 0 (independent stimuli) to 0.99.

### Generating ensembles of networks

For the all excitatory and all inhibitory networks of 4 neurons, we simulated the all 4096 unweighted and directed topologies. For networks of 15 neurons, the number of different unweighted directed topologies is larger than 10^63^. We therefore estimated the similarity metrics for an ensemble of 10,000 unweighted directed networks, sampled from an Erdős—Renyi random graph model [12], where the probability of forming a synapse between any two neurons in a network was *p_syn_* = 0.5. Each of the 15 neurons was randomly assigned a “type” - excitatory or inhibitory - with the probability of a random cell being inhibitory was *p_inhib_* = 0.2. This determined the sign of each synapse in the network. Synaptic weights were drawn from a log-normal distribution *σ =* 0.5 [42]. The mean strength of inhibitory synapses was scaled to ensure *p_inhib_*〈*w_inhib_*〉 = (1 − *p_inhib_*) 〈*w_ex_*〉, where the average is over the entire ensemble of networks. Networks of 50 or 100 neurons were randomly generated in a similar manner such that the probability of forming a synapse between two arbitrary neurons is *p_syn_* = 0.1, and *p_inhib_* = 0.2. (*p_syn_* = 0.5 and *p_syn_* = 0.05 were also considered and gave similar results).

### Permutation of synapses for networks of 50 and 100 neurons

The original random networks (see main text) were shuffled to generate variants that retained the total sum of synaptic input to the i-th neuron, by permuting all 49 (or 99) off-diagonal elements of the i-th column in the network’s connectivity matrix. Variants that retained total sum of synaptic outputs were similarly generated by permuting off-diagonal elements of each rows. Control networks were generated by shuffling all off-diagonal elements, irrespective of their original row or column.

### Fitting pairwise maximum entropy models

For large networks, direct estimation of *P_emp_* (**x**|*s*) would require prohibitive or unrealistic long stimuli (since for a network of N neurons, the number of possible activity patterns is 2^*N*^). We therefore used pairwise maximum entropy models to describe the response distributions. These models have been shown to be highly accurate for the activity patterns of groups of this scale [10], and we validated that this was the case for our networks. For each network and stimulus, we learned the maximum entropy model of its responses to the stimulus based on the firing rates and pairwise correlations between cells, given by 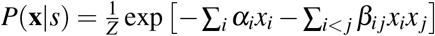; Models were learned using the MaxEnt toolbox software [54].

### Finding a sparse representation of structural features based on Cholesky decomposition

The Cholesky decomposition of a Hermitian positive-definite matrix [55] factorizes it uniquely into the product of a lower triangular matrix and its conjugate transpose. To interpret the structure of the matrix *M*\* based on the decomposition *M*\* = *R* · *R^T^*, we used the fact that right-multiplying *R* by any unitary matrix *U* results in a decomposition *M*\* = (*RU*)(*RU*)^*T*^, which means ||*R*^*T*^*g* − *R*^*T*^*g*′||^2^ = ||(*RU*)^*T*^*g* − (*RU*)^*T*^*g*′||^2^. We therefore solved the constrained optimization problem: *L* = argmin_*Uϵ{Q|QQ^T^*=*I}*_||*RU*||_1_ over the manifold of all possible unitary matrices [56, 57], and found a matrix *U* such that *L* = *RU* is maximally sparse and yet remains an exact decomposition of *M*.

### Multidimensional Scaling

Given an *N* × *N* dissimilarity matrix *D_func_* and a desired number of dimensions *p* (p=2 in the main text), we found the *N* × *p* embedding matrix such that the pairwise distances in the p-dimensional space minimize the mean squared error with respect to the dissimilarities specified by *D*. MDS was implemented using the scikit-learn python package [58].

### Divergences between probability distributions

The Kullback-Leibler divergence between probability distributions *P, Q* is defined as 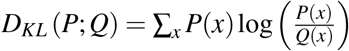, which measures the distinguishability of distributions in bits, and has multiple information theory and statistics motivations and interpretations [59]. The Jensen-Shannon divergence [60] is a symmetric and bounded extension of the Kullback-Leibler divergence, defined as 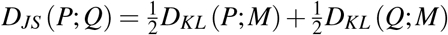 where 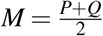. This is a measure of dissimilarity between probability distributions, which is 0 bits for identical distributions, and 1 bit for non-overlapping distributions.

## ACKNOWLEDGMENTS

We thank Roy Harpaz, Udi Karpas, Tal Tamir, Yoni Mayzel, Omri Camus, Benny Brazowski, Gasper Tkacik, and Allan Drummond for discussions, comments, and ideas. ES was supported by the European Research Council (ERC 311238 NEURO-POPCODE), Simons Collaboration on the Global Brain (542997), the Israel Science Foundation (Grant 1629/12), the Israel–US Binational Science Foundation, research support from Martin Kushner Schnur and Mr. and Mrs. Lawrence Feis, and is the Joseph and Bessie Feinberg Professorial Chair.

## SUPPLEMENTARY MATERIALS

Supplementary Text

Figs. S1 to S9

References (*61–67*)

## REFERENCES

1. Marder, E. Variability, compensation, and modulation in neurons and circuits. Proceedings of the National Academy of Sciences 108, 15542–15548 (2011).

2. Barkai, N. & Leibler, S. Robustness in simple biochemical networks. Nature 387, 913–917 (1997).

3. Tsai, T. Y.-C. et al. Robust, tunable biological sscillations from interlinked positive and negative feedback Loops. Science 321, 126–129 (2008).

4. Albert, R., Jeong, H. & Barabasi, A.-L. Error and attack tolerance of complex networks. Nature 406, 378–382 (2000).

5. Gu, S. et al. Controllability of structural brain networks. Nature Communications 6, 8414 (2015).

6. Atick, J. J. Could information theory provide an ecological theory of sensory processing? Network: Computation in Neural Systems 3, 213–251 (1992).

7. Tkačik, G., Prentice, J. S., Balasubramanian, V. & Schneidman, E. Optimal population coding by noisy spiking neurons. Proceedings of the National Academy of Sciences of the United States of America 107, 14419–14424 (2010).

8. Mora, T. & Bialek, W. Are Biological Systems Poised at Criticality? Journal of Statistical Physics 144, 268–302 (2011).

9. Rubinov, M., Sporns, O., Thivierge, J.-P. & Breakspear, M. Neurobiologically Realistic Determinants of Self-Organized Criticality in Networks of Spiking Neurons. PLoS Computational Biology 7, e1002038 (2011).

10. Ganmor, E., Segev, R. & Schneidman, E. Sparse low-order interaction network underlies a highly correlated and learnable neural population code. Proceedings of the National Academy of Sciences 108, 9679–9684 (2011).

11. Zurn, P. & Bassett, D. S. Network architectures supporting learnability. Philosophical Transactions of the Royal Society B: Biological Sciences 375, 20190323 (2020).

12. Newman, M. E. J. The Structure and Function of Complex Networks. SIAM Review 45, 167–256 (2003).

13. Milo, R. et al. Network motifs: simple building blocks of complex networks. Science 298, 824–827 (2002).

14. Barabási, A.-L. & Albert, R. Emergence of Scaling in Random Networks. Science 286, 509–512 (1999).

15. Bullmore, E. T., Sporns, O. & Solla, S. A. Complex brain networks: graph theoretical analysis of structural and functional systems. Nature Reviews Neuroscience 10, 186–98 (2009).

16. Olshausen, B. A. & Field, D. J. Sparse coding with an overcomplete basis set: A strategy employed by V1? Vision Research 37, 3311–3325 (1997).

17. Ganguli, S. & Sompolinsky, H. Compressed Sensing, Sparsity, and Dimensionality in Neuronal Information Processing and Data Analysis. Annual Review of Neuroscience 35, 485–508 (2012).

18. Maoz, O., Tkačik, G., Esteki, M. S., Kiani, R. & Schneidman, E. Learning probabilistic representations with randomly connected neural circuits. bioRxiv:478545 (2018).

19. Soriano, J., Martinez, M. R., Tlusty, T. & Moses, E. Development of input connections in neural cultures. Proceedings of the National Academy of Sciences 105, 13758–13763 (2008).

20. Ponce-Alvarez, A., Jouary, A., Privat, M., Deco, G. & Sumbre, G. Whole-Brain Neuronal Activity Displays Crackling Noise Dynamics. Neuron 100, 1446–1459 (2018).

21. Bargmann, C. I. & Marder, E. From the connectome to brain function. Nature Methods 10, 483–490 (2013).

22. Lichtman, J. W., Pfister, H. & Shavit, N. The big data challenges of connectomics. Nature Neuroscience 17, 1448–54 (2014).

23. Jiang, X. et al. Principles of connectivity among morphologically defined cell types in adult neocortex. Science 350, aac9462–aac9462. arXiv: 15334406 (2015).

24. Oh, S. W. et al. A mesoscale connectome of the mouse brain. Nature 508, 207–214 (2014).

25. Helmstaedter, M. et al. Connectomic reconstruction of the inner plexiform layer in the mouse retina. Nature 500, 168–174 (2013).

26. Xu, C. S. et al. A Connectome of the Adult Drosophila Central Brain. biorXiv:2020.01.21.911859 (2020).

27. Varshney, L. R., Chen, B. L., Paniagua, E., Hall, D. H. & Chklovskii, D. B. Structural Properties of the Caenorhabditis elegans Neuronal Network. PLoS Computational Biology 7, e1001066 (2011).

28. Prinz, A. A., Bucher, D. & Marder, E. Similar network activity from disparate circuit parameters. Nature neuroscience 7, 1345–52 (2004).

29. Jun, J. J. et al. Fully integrated silicon probes for high-density recording of neural activity. Nature 551, 232–236 (2017).

30. Susaki, E. A. et al. Whole-brain imaging with single-cell resolution using chemical cocktails and computational analysis. Cell 157, 726–39 (2014).

31. Cossell, L. et al. Functional organization of excitatory synaptic strength in primary visual cortex. Nature 518, 399–403 (2015).

32. Scala, F. et al. Layer 4 of mouse neocortex differs in cell types and circuit organization between sensory areas. Nature Communications 10, 4174 (2019).

33. Wanner, A. A. & Friedrich, R. W. Whitening of odor representations by the wiring diagram of the olfactory bulb. Nature Neuroscience 23, 433–442 (2020).

34. Litwin-Kumar, A. & Turaga, S. C. Constraining computational models using electron microscopy wiring diagrams. Current Opinion in Neurobiology 58, 94–100 (2019).

35. Sizemore, A., Giusti, C., Betzel, R. F. & Bassett, D. S. Closures and Cavities in the Human Connectome. arXiv:1608.03520. arXiv: 1608.03520 (2016).

36. Timme, M. & Casadiego, J. Revealing networks from dynamics: an introduction. Journal of Physics A: Mathematical and Theoretical 47, 343001 (2014).

37. Tikhonov, M. & Bialek, W. Complexity of generic biochemical circuits: topology versus strength of interactions. Physical Biology 13, 066012 (2016).

38. Morrison, K. & Curto, C. in Algebraic and Combinatorial Computational Biology 241–277 (Elsevier, 2019).

39. Mastrogiuseppe, F. & Ostojic, S. Linking Connectivity, Dynamics, and Computations in Low-Rank Recurrent Neural Networks. Neuron 99, 609–623.e29 (2018).

40. Raman, D. V., Rotondo, A. P. & O’Leary, T. Fundamental bounds on learning performance in neural circuits. Proceedings of the National Academy of Sciences 116, 10537–10546 (2019).

41. Brunel, N. Dynamics of Sparsely Conntected Networks of Excitatory and Inhibitory Spiking Neurons. Journal of Computational Neuroscience 8, 183–208 (2000).

42. Buzsáki, G. & Mizuseki, K. The log-dynamic brain: how skewed distributions affect network operations. Nature Reviews Neuroscience 15, 264–78 (2014).

43. Kulis, B. Metric Learning: A Survey. Foundations and Trends in Machine Learning 5, 287–364 (2013).

44. Perrot, M., Habrard, A., Muselet, D. & Sebban, M. in Computer Vision - ECCV 2014 96–111 (Springer International Publishing, 2014).

45. Freeman, J. & Simoncelli, E. P. Metamers of the ventral stream. Nature Neuroscience 14, 1195–1201 (2011).

46. Ganmor, E., Segev, R. & Schneidman, E. A thesaurus for a neural population code. eLife 4, e06134 (2015).

47. Chaudhuri, R., Gerçek, B., Pandey, B., Peyrache, A. & Fiete, I. The intrinsic attractor manifold and population dynamics of a canonical cognitive circuit across waking and sleep. Nature Neuroscience 22, 1512–1520 (2019).

48. Sizemore, A. E. et al. Cliques and cavities in the human connectome. Journal of Computational Neuroscience 44, 115–145 (2017).

49. Van Nimwegen, E., Crutchfield, J. P. & Huynen, M. Neutral evolution of mutational robustness. Proceedings of the National Academy of Sciences 96, 9716–9720 (1999).

50. Turrigiano, G. G. & Nelson, S. B. Homeostatic Plasticity in the Developing Nervous System. Nature Reviews Neuroscience 5, 97–107 (2004).

51. Royer, S. & Pare, D. Conservation of total synaptic weight through balanced synaptic depression and potentiation. Nature 422, 518–522 (2003).

52. Linssen, C. et al. NEST 2.16.0. Version 2.16.0. doi.org/10.5281/zenodo.1400175 (2018).

53. Kuhn, A., Aertsen, A. & Rotter, S. Higher-Order Statistics of Input Ensembles and the Response of Simple Model Neurons. Neural Computation 15, 67–101 (2003).

54. Maoz, O. & Schneidman, E. maxent_toolbox: Maximum Entropy Toolbox for MATLAB, version 1.0.2. Version 1.02. doi:10.5281/zenodo.191625 (2017).

55. Nash, J. C. Compact numerical methods for computers: linear algebra and function minimisation eng;eng (Hilger, Bristol, 1979).

56. Absil, P.-A., Mahony, R. & Sepulchre, R. Optimization Algorithms on Matrix Manifolds (Princeton University Press, USA, 2007).

57. Townsend, J., Koep, N. & Weichwald, S. Pymaopt: A Python Toolbox for Optimization on Manifolds using Automatic Differentiation. Journal of Machine Learning Research 17, 1–5 (2016).

58. Pedregosa, F. et al. Scikit-learn: Machine Learning in Python. Journal of Machine Learning Research 12, 2825–2830 (2011).

59. MacKay, D. J. C. Information Theory, Inference & Learning Algorithms (Cambridge University Press, New York, NY, USA, 2002).

60. Lin, J. Divergence measures based on the Shannon entropy. IEEE Trans. Information Theory 37, 145–151 (1991).

61. Rotter, S. & Diesmann, M. Exact digital simulation of time-invariant linear systems with applications to neuronal modeling. Biological Cybernetics 81, 381–402 (1999).

62. Burkitt, A. N. A Review of the Integrate-and-fire Neuron Model: I. Homogeneous Synaptic Input. Biological Cybernetics 95, 1–19 (2006).

63. Stimberg, M., Brette, R. & Goodman, D. F. Brian 2, an intuitive and efficient neural simulator. eLife 8, e47314 (2019).

64. Stimberg, M., Goodman, D. F. M., Brette, R. & De Pitta, M. Modeling neuron-glia interactions with the Brian 2 simulator. bioRxiv:198366 (2017).

65. Virtanen, P. et al. SciPy 1.0: fundamental algorithms for scientific computing in Python. Nature Methods 17, 261–272 (2020).

66. Hagberg, A. A., Schult, D. A. & Swart, P. J. Exploring Network Structure, Dynamics, and Function using NetworkX in Proceedings of the 7th Python in Science Conference (eds Varoquaux, G., Vaught, T. & Millman, J.) (Pasadena, CA USA, 2008), 11–15.

67. Wills, P. & Meyer, F. G. Metrics for Graph Comparison: A Practitioner’s Guide 2020.

