## Supplementary Materials for "Learning the architectural features that predict functional similarity of neural networks"

### SUPPLEMENTARY TEXT

**Identifying stimulus strengths that elicit networks' responses.** We repeated the analysis of network responses and similarity for a wide range of external input rates - from very weak stimuli that elicited almost no spiking responses to very strong stimuli that elicited firing rates that saturated our chosen binning resolution. The relation between the firing rates and the stimulus strength parameter $\eta$, for the ensemble of 15 neurons is shown in Fig S1, and was used to set the range of stimulus values that were studied in detail along the manuscript; The networks analyzed in Fig. 1-3 in the main text use $\eta = 1.5$.

**Different choice for temporal bin size, synaptic dynamics, and single neuron models.** To verify our results did not depend on a specific choice of the simulation configuration, we repeated the analysis in the main text for different simulation and analysis parameters:

1. *Temporal bin sizes.* The analysis presented in main text used time bin of $\Delta t = 20ms$. We have repeated the analysis with $\Delta t = 10ms$ and $40ms$.

2. *Synaptic dynamics.* Throughout the main text, the synaptic dynamics of the simulated networks followed an alpha-function, $I(t) = I_0 \cdot \frac{t}{\tau} \cdot e^{1-\frac{t}{\tau}}$ [61], in which post-synaptic potentials have a finite rise time. We have repeated the analysis with delta-function activation function of synapses [62], in which post-synaptic potential jumps on each spike arrival.

3. *Neuron model.* The neuron model used in the main text was a current-based model ("iaf psc alpha" in NEST [52]), in which sub-threshold dynamics are linear, and threshold crossing is followed by an absolute refractory period. Conductance based networks were simulated using Brian 2 [63], and simulation parameters were adopted from [64].

Similar to the results presented in the main text, the model based on the full Mahalanobis matrix, the complete feature-based model, and the model based on the total synaptic inputs, all showed significantly higher correlation with $D_{func}$ compared to others structural measures; Euclidean distance is shown in figure S2 as an example, but the behavior for other structural metrics is similar.

**Predicting PSTH-based functional similarity.** The similarity between the spiking responses of networks can be evaluated using different metrics, and the specific choice of a metric highlights different aspects of the neural code. It is not immediately clear whether the model defined in equation 3, which accurately predicts $D_{func}$ (as defined in the main text), would generalize to other functional metrics. We therefore asked whether our feature-based model can predict a PSTH-based similarity measure of the neural responses.

We used the binarized spiking response of network $G$ to stimulus $s$ (a matrix of the form $\{0,1\}^{N\times T}$ as in Fig. 1), which we denote by $x_G(s)$, to get the time-dependent firing rate or post stimulus time histogram (PSTH): For a given stimulus $s$ and for each network $G$, we convolve the i-th row of $x_G(s)$ with a 200 ms sliding window to get the PSTH of the i-th neuron in network $G$, $r^i_G(t)$ (which is a vector of real number of size $T-W$). We then measure the PSTH-based functional dissimilarity between two networks as the average over neurons of the PSTH difference between the corresponding neurons in each network:

$$D_{PSTH}\left(G, G'|s\right) = \frac{1}{N}\sum_{i=1}^{N} \|r^i_G(t) - r^i_{G'}(t)\|_2$$

We found that $D_{func}$ and $D_{PSTH}$ are highly correlated across different stimuli. Consequently, our feature based model accurately predicts the PSTH-based functional dissimilarity, as shown in Figure S3.

**Low-dimensional complexity of the dissimilarity matrix between networks.** To assess the lower dimensional structure of the dissimilarity matrix, we compared its spectrum to that of randomly shuffled controls. In Fig. S4 we show for three representative example networks of size 15 responding to three different stimuli, that the spectrum of the original dissimilarity matrix ("empirical" in blue) decays significantly faster than that of randomly shuffled version of these matrices ("control" in black). Thus, the lower dimensional structure of these matrices depends on the relations between all the pairwise distances and not merely their overall distribution.

**Comparing the predictive power of different common structural metrics.** We explored a wide range of structural metrics and asked how well they predicted the functional dissimilarity of network responses. Figure S5 shows an extended version of figure 2 in the main text: accuracy was measured by the correlation between the distances computed by each of the methods and the functional dissimilarity $D_{func}$. As in figure 2, the stimulus to each neuron was an independent realization of the same Poisson inputs, with $N = 15, \eta = 1.5$ and $\rho = 0$. Our learned model (black) outperformed all metrics by a large margin.

Details of the metrics used in the comparison:

1. Metrics between continuous vectors:
   - $L_1$ (Manhattan) - $\sum|x_i - y_i|$
   - Braycurtis - $\sum_i \frac{|x_i - y_i|}{|x_i + y_i|}$
   - Canberra - $\sum_i \frac{|x_i - y_i|}{|x_i| + |y_i|}$
   - $L_\infty$ (Chebyshev) - $\max|x_i - y_i|$
2. Metrics between binary vectors (applied to binarized connectivity matrix):
   - Dice - $\frac{\sum|x_i - y_i|}{2\sum x_i \cdot y_i + \sum|x_i - y_i|}$
   - Jaccard - $\frac{\sum|x_i - y_i|}{\sum x_i \cdot y_i + \sum|x_i - y_i|}$
   - Kulsinki - $\frac{\sum|x_i - y_i| - \sum x_i \cdot y_i + N(N-1)}{\sum|x_i - y_i| + N(N-1)}$
   - Rogers-Tanimoto - $\frac{2\sum|x_i - y_i|}{\sum x_i \cdot y_i + \sum(x_i - 1)\cdot(y_i - 1) + 2\sum|x_i - y_i|}$
   - Russel-Rao - $\frac{N(N-1) - \sum x_i \cdot y_i}{N(N-1)}$
   - Sokal-Sneath - $\frac{2\sum|x_i - y_i|}{\sum x_i \cdot y_i + 2\sum|x_i - y_i|}$
   - Yule - $\frac{2\sum|x_i - y_i|}{\sum x_i \cdot y_i \cdot \sum(x_i - 1)\cdot(y_i - 1) + \sum|x_i - y_i|}$

All structural metrics were computed using the SciPy python package [65]. Spectral distances were computed as follows: For each connectivity matrix $G$, the directed graph Laplacian was computed using 3 different algorithms, as implemented by the networkx python package [66]. The real part of each Laplacian spectrum was computed and sorted, yielding a vector of size $N$. Finally, Euclidean pairwise distances were computed between the vectors [67].

**Finding the optimal Mahalanobis matrix $M^*$ under a regularized optimization.** The optimal Mahalanobis matrix $M^*$ was found by solving the optimization problem described in the main text (equation 2). For networks of size $N$, the number of parameters in $M^*$ is $N^2 \cdot (N-1)^2$. To avoid overfitting, we used a regularization term that was weighted by a free parameter $\alpha$. For networks of size 15 we solved the optimization problem with 50 different values of $\alpha$, equally spaced on a logarithmic scale from $10^{-6}$ to $10^6$. For each value of $\alpha$ we estimated the accuracy of the resulting $M^*$ on a held-out validation set (a quarter of the size of the train set). The value of $\alpha$ that gave the minimal loss for each stimulus was used to obtain $M^*$ that were used in the main text; with some small variation across stimuli, $\alpha_{opt} \approx 10^3$.

**The optimal Mahalanobis $M^*$ for shuffled data shows no structure.** To verify that the success and accuracy of our learned Mahalanobis matrix $M^*$ was not a result of overfitting or of over-expressive power of this model, we repeated the fitting procedure with shuffled functional dissimilarity values, by randomly permuting the entries of the dissimilarity matrix. For different values of the regularization parameter $\alpha$ we solved the optimization problem and assessed predictive performance on a validation set, as described above. The value of $\alpha$ that gave the best performance on the validation set was used to compute $M^*$, and its performance was measured on a test set, shown in Fig. S7. Unlike the case for the real data, the optimal $M^*$ was unable to model the shuffled data. This result, together with the cross-validation strategy described above, show that the predictive accuracy of $M^*$ stems from its ability to capture the geometry of the functional space of networks, and not from its computational expressive power.

**Correlation between dissimilarity matrices across the space of stimuli.** The functional dissimilarity matrix between networks depends on the stimulus that is presented to the networks. It is not immediately clear that these dissimilarity matrices would be related in a simple way. In particular, networks that respond very differently to one stimulus may have very similar responses to another stimulus. To investigate this, we calculated the Pearson correlation between all computed dissimilarity matrices for the different stimuli, and the one presented in the main text (for which $\eta = 1.5$ and $\rho = 0$).

Figure S8 shows that over large parts of the space of stimuli, functional dissimilarity matrices were highly correlated. This implies that the functional distances between networks were scaled on average by a multiplicative constant that was a function of the statistics of the stimulus. Up to this scaling behavior, the overall structure of functional networks space remained stable.

**Relative importance of the sum of synaptic inputs and the sum of synaptic outputs for the feature based models.** The models based on sum of synaptic inputs and sum of synaptic outputs (IO-based models), described by equation 3 in the main text, were fitted by a linear regression that predicts $D_{func}$ using a weighted sum of the squared differences between the sum of synaptic inputs and sum of synaptic outputs of each of the neurons. To characterize the relative importance of the synaptic inputs and synaptic outputs in predicting functional dissimilarity of networks, we fitted two additional models for each $(\eta, \rho)$ stimulus: A model that relies only on the sum of synaptic inputs of each neuron and a model that relies only on the sum of synaptic outputs of each neuron. We computed the accuracy of each model, defined as Pearson's correlation with empirical $D_{func}$, denoting the accuracy of the inputs-based model as $A_I$, and the accuracy of the outputs-based model as $A_O$. We then compared the accuracy of these two models by computing the ratio between correlation coefficients, $\frac{A_I}{A_O}$, as plotted in Fig. S9. Interestingly, we found a transition from an outputs-dominated to an inputs-dominated regime as stimulus strength increased. We note that the model in the main text used both the inputs and outputs as features, and therefore outperforms both, by construction.

### SUPPLEMENTARY FIGURES

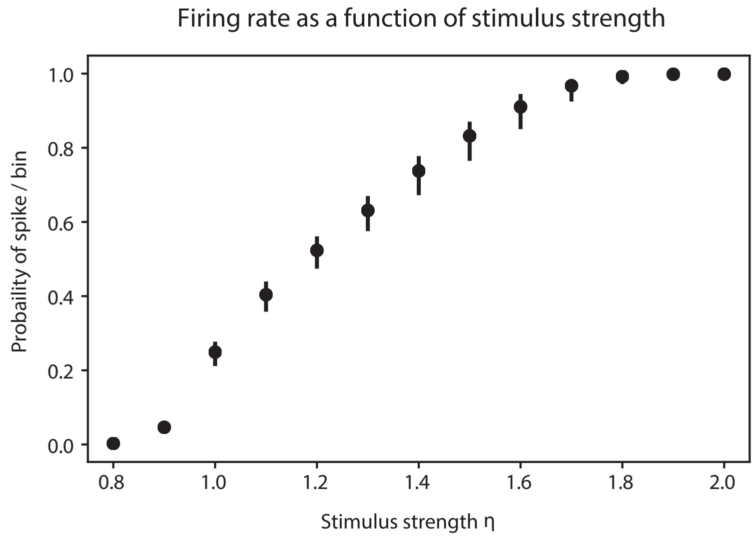


**Figure S1. Firing rates as a function of stimulus strength.** In the main text, results for the N=15 ensemble are for $\eta = 1.5$ unless stated otherwise.

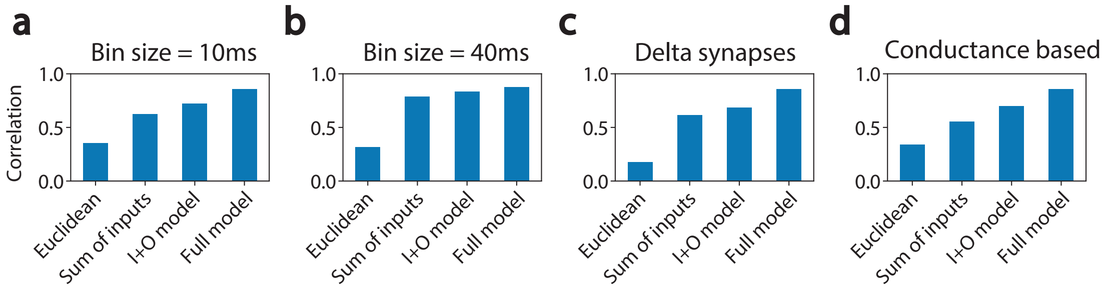


**Figure S2. Comparison of the performance of models under different neuron model, synaptic dynamics, and time bins.** Our model outperformed the Euclidean structural distance for different choice for temporal bin size (a,b), synaptic dynamics (c), and single neuron models (d).

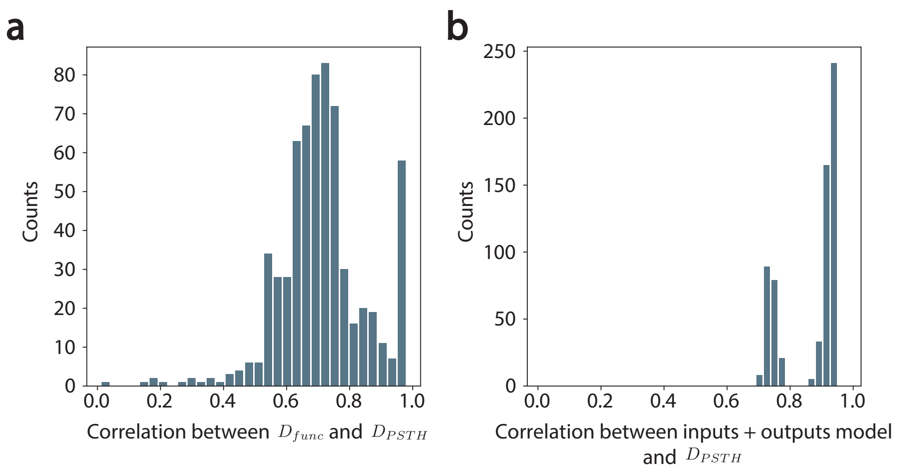


**Figure S3. Feature based model accurately predicts other functional dissimilarity measures. (a)** For different values of stimulus strength and correlation, the $D_{func}$ measure and $D_{PSTH}$ are highly correlated. **(b)** Similar to the results presented in the main text, a feature-based model that relies on the sum of synaptic inputs and sum of synaptic outputs of each neuron accurately predicts $D_{PSTH}$.

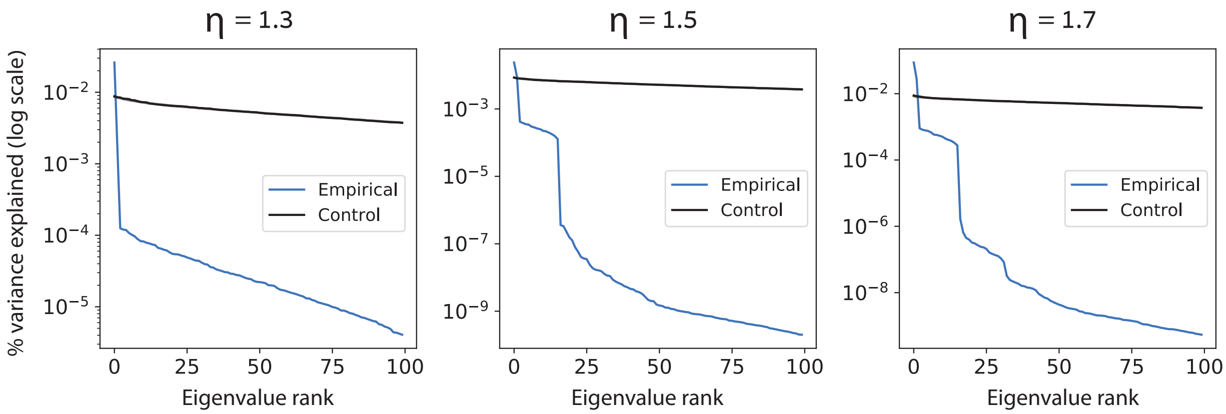


**Figure S4. Low dimensional structure of the matrix of distances between networks.** For 3 different stimuli, the spectrum of the empirical functional dissimilarity matrix decays faster than shuffled control.

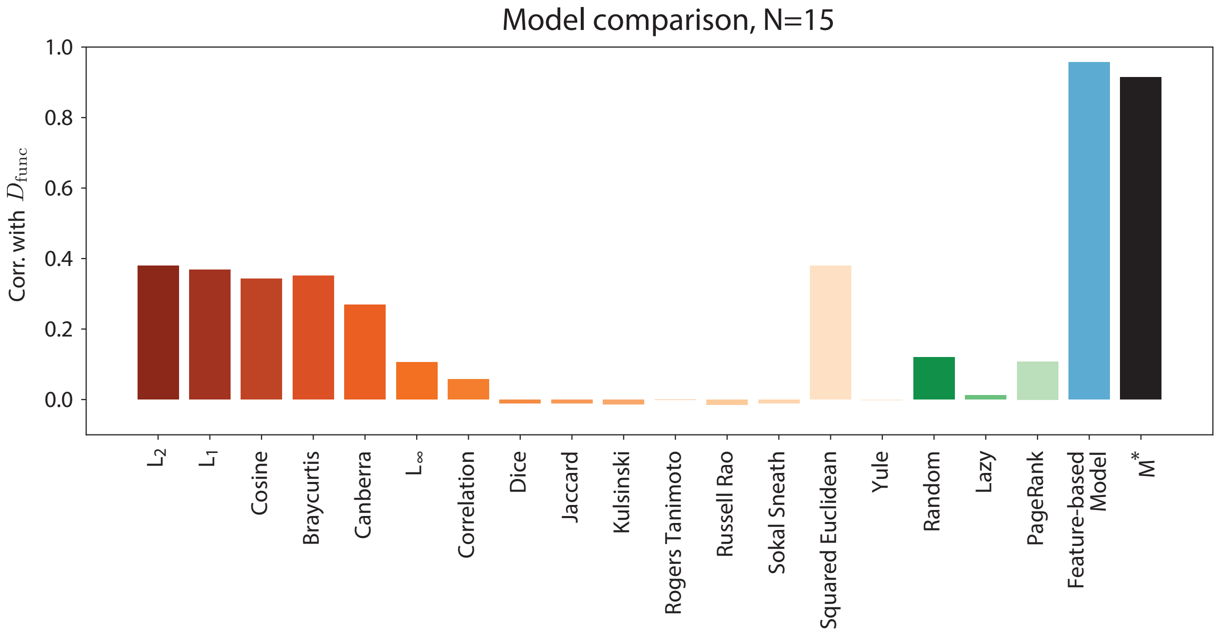


**Figure S5. A wide range of structural metrics fail to predict functional dissimilarity.** We compared a wide variety of structural metrics. The model based on $M^*$ (black) significantly outperformed all metrics we have considered. Results for the $N = 15$ ensemble and $\eta = 1.5$.

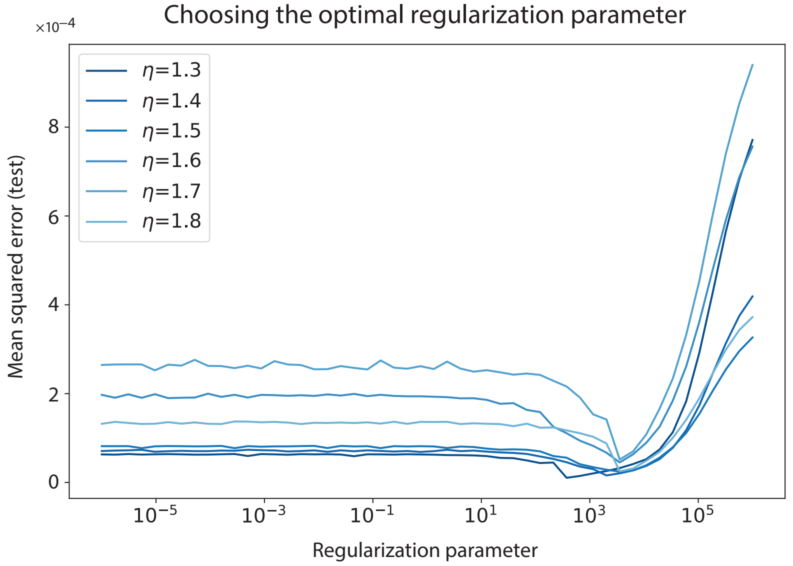


**Figure S6. Choosing the optimal value of the regularization parameter $\alpha$ using train-test splitting.** For the ensemble of networks with $N = 15$, $M^*$ was found with regularized optimization for 50 different values of $\alpha$.

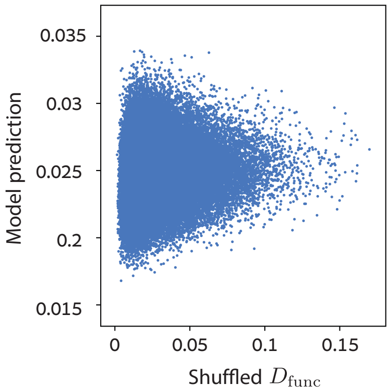


**Figure S7. Fitting $M^*$ to shuffled data.** The optimal Mahalanobis $M^*$ shows no structure.

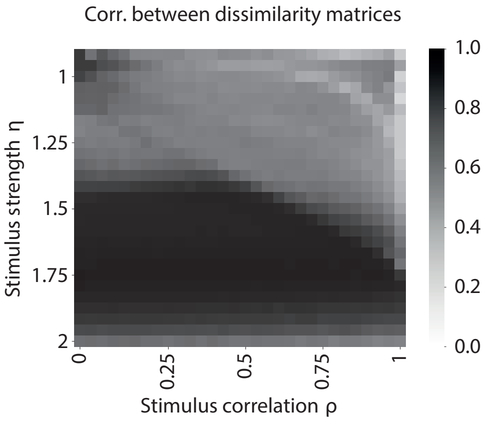


**Figure S8. Correlation between dissimilarity matrices across the space of stimuli.** Mean correlation across stimulus space is 0.8 with a standard deviation of 0.16 (median 0.81).

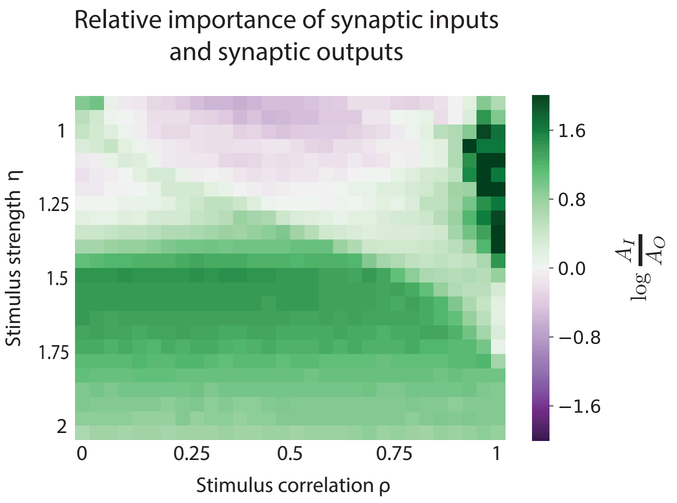


**Figure S9. Relative importance of the sum of synaptic inputs and the sum of synaptic outputs for the feature based models varies across stimulus space**. As stimuli strength increases, similarity of synpatic inputs dominates similarity of synaptic outputs.
